# Ancient pyroptotic machinery via GSDMA/B cleavage by LPS-activated caspase-1 in cartilaginous fish

**DOI:** 10.64898/2026.07.03.736391

**Authors:** Xuxia Wei, Ru Zhuang, Xueqin Jia, Xinli Wang, Suizhi Li, Ziwen Huang, Gaijing Zhou, Anlong Xu, Shaochun Yuan

**Affiliations:** Guangdong Province Key Laboratory of Pharmaceutical Functional Genes, MOE Key Laboratory of Gene Function and Regulation, MOE Engineering Center of South China Sea Marine Biotechnology, State Key Laboratory of Biocontrol, School of Life Sciences, Sun Yat-Sen University, Guangzhou 510275, China; Laboratory for Marine Biology and Biotechnology, Pilot National Laboratory for Marine Science and Technology, Qingdao 266237, China; Sun Yat-sen University Institute of Advanced Studies, Hong Kong SAR 999077, China

**Keywords:** Pyroptosis, gasdermin, caspase-1, *Callorhinchus milii*, non-canonical inflammasome

## Abstract

Gasdermins are pore-forming effectors that mediate pyroptosis, an inflammatory form of programmed cell death characterized by membrane permeabilization and the release of intracellular contents. Phylogenetically, gasdermin members can be broadly divided into two major branches, the GSDME/PJVK branch and the GSDMA/B/C/D branch. Whereas the GSDME mediated pyroptosis was traced back to metazoans, the functional origins of GSDMA/B/C/D branch remain poorly understood. As a basal representative of the GSDMA-D lineage in cartilaginous fish, *Callorhinchus milii* GSDMA/B (CmiGSDMA/B) provides essential information for the ancestral state of this branch. Here, we functionally characterized CmiGSDMA/B and identified CmiCASP1 as its upstream protease. Mechanistically, Lipopolysaccharide (LPS) activates CmiCASP1 via its CARD domain, leading to cleavage of CmiGSDMA/B into two functionally distinct products, N241 and N288. N241 binds cell membrane to drive pyroptosis, whereas N288 suppresses N241-triggered cell death. Interestingly, N241 exhibits bactericidal activity against Gram-negative bacteria in vitro, suggesting that antimicrobial activity may have been an early feature of the GSDMA-D lineage. Collectively, these findings provide insight into a non-canonical, LPS-responsive, caspase-driven pyroptosis pathway in cartilaginous fish and reveal the dual-fragment antagonistic regulation within this branch.

## Introduction

Pyroptosis is a lytic and proinflammatory form of programmed cell death that plays a critical role in host defense and innate immunity (Broz, 2025; Cookson and Brennan, 2001). It is executed by pore-forming gasdermin (GSDM) family members in response to extracellular or intracellular injury cues (Broz *et al*., 2020). GSDMs typically consist of a pore-forming N-terminal domain (NTD) and an autoinhibitory C-terminal domain (CTD) linked by a flexible linker (Broz *et al*., 2020; Ding *et al*., 2016). In the resting state, GSDMs are maintained in an autoinhibitory conformation via intramolecular interactions (Bai *et al*., 2025). Proteolytic cleavage within the interdomain linker releases the pore-forming NTD, which subsequently oligomerizes and inserts into the plasma membrane as large β-barrel pores, ranging from 15-21.5 nm (Ruan *et al*., 2018; Wang *et al*., 2023a; Xia *et al*., 2021). Permeabilization of the plasma membrane disrupts cellular integrity, drives cellular swelling and results in the release of danger-associated molecular patterns (DAMPs) and cytokines, such as IL-1β and IL-18 (Shi *et al*., 2017). Pyroptosis serves as a critical antimicrobial defense mechanism, yet its dysregulated activation drives inflammatory pathogenesis and autoimmune diseases (Liu *et al*., 2021).

*GSDMs* and *GSDM*-like genes constitute an evolutionarily conserved protein family distributed across metazoans, fungi, bacteria, and viruses (Boys *et al*., 2023; Daskalov *et al*., 2020; Johnson *et al*., 2022; Yuan *et al*., 2022). Microbial GSDM-like homologs retain pore-forming capacity and can be activated by caspase-like proteases, indicating that proteolytic relief of C-terminal autoinhibition represents an ancestral immune strategy predating metazoan evolution (Clavé *et al*., 2022; Johnson *et al*., 2022). Metazoan GSDMs can be divided into two major clades, the GSDME/PJVK clade and the GSDMA/B/C/D clade (Angosto-Bazarra *et al*., 2022; Wang *et al*., 2023b). GSDME represents the most ancestral metazoan gasdermin, displaying broad phylogenetic distribution from cnidarians to mammals and executing evolutionarily conserved pyroptotic functions (Wang *et al*., 2023b). For instance, human GSDME functions as a molecular switch that converts non-inflammatory apoptosis into inflammatory pyroptosis upon caspase-3 mediated cleavage at Asp270 in cancer cells (Wang *et al*., 2017), while caspase-3/GSDME axis in invertebrate phyla primarily orchestrates innate immune defense responses, such as cnidarian and lancelet GSDME homologs (Jiang *et al*., 2020; Qin *et al*., 2023). Notably, proteolytic diversification in lancelet GSDME yields two N-terminal variants with distinct membrane-permeabilizing capacities (Wang *et al*., 2023b), adding regulatory sophistication to GSDME-mediated membrane permeabilization.

Unlike GSDME/PJVK clade which spans the animal kingdom, the GSDMA/B/C/D clade is restricted to jawed vertebrates. Mammalian genomes encode multiple paralogs (GSDMA-D), differing in tissue distribution, proteolytic activation mechanisms, and physiological or pathological outcomes (Zhu *et al*., 2024). GSDMD, the most extensively studied member, functions as a central executioner of pyroptosis when activated canonically by caspase-1 downstream of inflammasome assembly, or non-canonically by caspase-4/5/11 upon cytosolic LPS sensing (Kayagaki *et al*., 2015; Lee *et al*., 2018; Shi *et al*., 2015). In contrast to the ubiquitous expression of GSDMD, GSDMA-C exhibit tissue-restricted distribution and mediate pathogen driven pathology in specific organs (Zhu *et al*., 2024). For instance, human GSDMA is highly expressed in skin epithelial cells and can be activated by skin-associated bacterial proteases, including SpeB and ScpA (Deng *et al*., 2022; LaRock *et al*., 2022; LaRock *et al*., 2025). GSDMC is predominantly expressed in intestinal epithelial cells and is implicated in type 2 immunity against intestinal parasites (Xi *et al*., 2021; Zhao *et al*., 2022). GSDMB is mainly expressed in gastric and respiratory epithelium and represents the most significant asthma susceptibility gene within the GSDM family (Das *et al*., 2016). Besides functional diversification, mammalian GSDMA and GSDMC display lineage-specific tandem gene expansions, as illustrated by the murine *Gsdma1-3* and *Gsdmc1-4* clusters (Qiu *et al*., 2017), indicative of diversifying selection driving their adaptive radiation and neofunctionalization. Recent studies have also revealed that non-mammalian GSDMA (in birds, reptiles, and amphibians) retains the caspase-1 cleavage site, whereas mammalian GSDMA has lost this site with its function being supplanted by GSDMD (Billman *et al*., 2024). Thus, the evolution from an ancestral pyroptosis executioner into multifunctional regulators drives the diversification and neofunctionalization of GSDMA-D paralogs involved in mucosal immunity.

Chondrichthyes (cartilaginous fish) constitute the most basal extant jawed vertebrates, offering unique insights into the ancestral state of vertebrate evolution (Venkatesh *et al*., 2014). Previously, cartilaginous fish GSDMA/B was proposed to represent an ancestral-like member of vertebrate GSDMA-D lineage (Wang *et al*., 2023b). In this study, by focusing on *Callorhinchus milii* GSDMA/B (CmiGSDMA/B) and its upstream inflammatory caspase, CmiCASP1, we found that LPS directly engages the CARD domain of CmiCASP1, triggering its activation, which subsequently promotes the proteolytic cleavage of CmiGSDMA/B, yielding two N-terminal fragments with opposite functions. Consistent with GSDMD, the functional N241 of CmiGSDMA/B can mediate pyroptosis and exhibit bactericidal activity against Gram-negative bacteria in vitro. Thus, our findings uncover an ancestral LPS-sensing CASP1-GSDMA/B axis in cartilaginous fish, revealing the pre-tetrapod state of this pore-forming family and illuminating the evolutionary transition from primitive cytolysis to sophisticated mucosal immunity.

## Results

### CmiCASP1-mediated cleavage activates CmiGSDMA/B to execute pyroptosis in the elephant shark (*Callorhinchus milii*)

Based on the obtained sequences by searching against publicly available genomic, EST, and protein databases, we re-constructed a maximum likelihood (ML) tree of GSDM family using bacterial and fungal GSDM-like homologs as outgroups (**Figure 1—figure supplement 1A**). As previously demonstrated, GSDMA/B sequences from basal jawed vertebrates (represented by *Callorhinchus milii* and *Rhincodon typus*) reside at the root of the GSDMA-D lineage (**Figure 1A** and **Figure 1—figure supplement 1A**) (De Schutter *et al*., 2021; Wang *et al*., 2023b). Moreover, as indicated in **Figure 1B**, CmiGSDMA/B retains the canonical domain organization of vertebrate GSDMs, including an N-terminal β-barrel pore-forming domain, a proteolytic linker region, and a C-terminal repressive domain. To identify potential activators of CmiGSDMA/B, we performed a genome-wide survey of caspases in *Callorhinchus milii*, revealing seven family members including CmiCASP1, a CARD-containing inflammatory caspase (**Figure 1—figure supplement 1B**). ML tree constructed based on the CASc (caspase catalytic) domains indicated that CmiCASP1 clusters with inflammatory caspases and occupies a basal position relative to tetrapod inflammatory caspases **(Figure 1C)**. Consistently, full-length sequence alignment also indicates that CmiCASP1 is more similar to tetrapod inflammatory caspases than to other caspase subfamilies (**Figure 1—figure supplement 1C**). Since caspases can be activated by autocleavage into the p20/p10 form (Lavrik *et al*., 2005), we predicted the p20/p10 form of CmiCASP1 and assessed its structural similarity to human caspases using RMSD and TM-score, where lower RMSD and higher TM-score values indicate greater structural similarity. Analyses showed that the active form of CmiCASP1 is much more structurally similar to HsaCASP1 than to HsaCASP3 (**Figure 1D**). Caspases typically recognize a tetrapeptide motif on the N-terminal side of the cleavage site, designated P4-P1, with cleavage occurring after the P1 residue (Talanian *et al*., 1997; Thornberry *et al*., 1997). These substrate residues are accommodated by four corresponding specificity pockets in the catalytic groove, S4-S1 (Thornberry *et al*., 1997). Comparative analysis of the catalytic domain showed that CmiCASP1 retains residues corresponding to the substrate-binding subsites of HsaCASP1, including residues contributing to the S1-S4 pockets (**Figure 1E** and **Figure 1—figure supplement 1D**). Among these pockets, the S4 pocket is especially important for distinguishing substrate preference because it accommodates the P4 residues, a major determinant of cleavage-site specificity (Nicholson, 1999; Poreba *et al*., 2013). Notably, the residue corresponding to HsaCASP1 Val348, located at the base of the S4 subsite that contributes to P4 substrate recognition, was conserved as a small hydrophobic residue in CmiCASP1 and other caspase-1 homologs, whereas the equivalent position in apoptotic caspase-3 is replaced by a bulky tryptophan residue (Wei *et al*., 2000). Structural modeling suggested that CmiCASP1 contains a relatively spacious and hydrophobic putative S4 pocket, resembling that of HsaCASP1 rather than the constricted and hydrophilic architecture of apoptotic caspase-3 (**Figure 1F**) (Poreba *et al*., 2013). In addition, sequence alignment identified C279 as the putative catalytic cysteine of CmiCASP1, corresponding to the conserved active-site cysteine of representative inflammatory caspases (**Figure 1—figure supplement 1D**). These sequence and structural features support the possibility that CmiCASP1 may act as an inflammatory caspase and nominate it as a candidate protease for CmiGSDMA/B processing.

**Figure 1.**
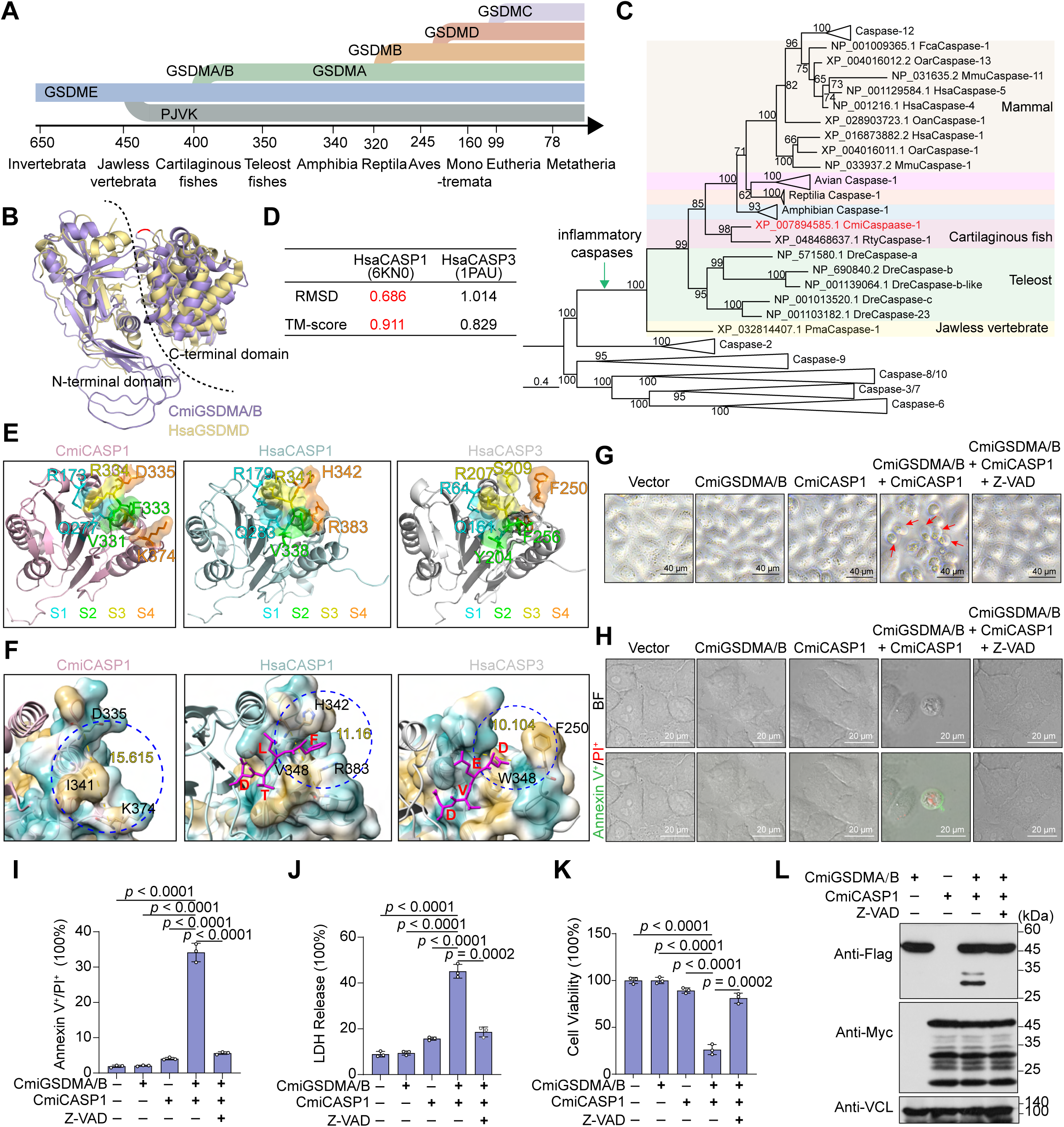
CmiCASP1 cleaves CmiGSDMA/B to execute pyroptosis in *C. milii*. (**A**) Timeline of GSDM gene family evolution showing phylogenetic branching in million years ago (MYA), the origin of ancestral GSDM, gene duplications giving rise to GSDME and PJVK, and lineage-specific expansions of GSDMA, GSDMB, GSDMC, and GSDMD in vertebrates. (**B**) Structural comparison of CmiGSDMA/B and HsaGSDMD. The predicted structure of CmiGSDMA/B (purple) was superimposed with the structure of HsaGSDMD (PDB: 6N9O) (yellow) using PyMOL. The N- and C-terminal domains are indicated and separated by a dashed line. (**C**) Phylogenetic analysis of caspases. Accession numbers are listed in **Supplementary file 1**. (**D**) RMSD and TM-score for structural comparisons of CmiCASP1 with HsaCASP1 and HsaCASP3. RMSD was calculated using PyMOL, and TM-score was calculated using Foldseek. (**E**) The distinct substrate-recognition pockets are shown in different colors, with the residues forming each pocket colored accordingly. (**F**) Hydrophobicity analysis of the S4 pocket. The region outlined by the blue dashed line corresponds to the S4 pocket. Cyan indicates hydrophilic regions, whereas yellow indicates hydrophobic regions. The magenta sticks indicate bound peptide ligands: FLTD in HsaCASP1 and DEVD in HsaCASP3. Residues forming the S4 pocket are labeled in black. The yellow label indicates the maximum distance accommodated by the S4 pocket. (**G**) Bright-field images of HeLa*^gsdmd/e DKO^* cells transfected with CmiGSDMA/B, CmiCASP1 or both, with or without Z-VAD-FMK treatment. Red arrows mark swollen, bubbling pyroptotic cells. (**H**) Microscopic analysis of cell death phenotypes using Annexin V-FITC/PI staining. Confocal microscopy images of HeLa*^gsdmd/e DKO^* cells transfected as indicated. (**I**) Flow cytometry analysis of Annexin V-FITC and PI double-stained cells. HEK293T cells were transfected and stimulated as indicated, followed by double staining with Annexin V-FITC and PI. (**J**) LDH release assay. LDH release was measured in the supernatants of HEK293T cells transfected with the indicated constructs. (**K**) ATP-based cell viability assay of HEK293T cells under different transfection and treatment conditions, as indicated. **(L)** Western blot analysis showing CmiCASP1-mediated cleavage of CmiGSDMA/B in HEK293T cells in the presence or absence of Z-VAD-FMK. Data are presented as mean ± SD from three independent experiments. The *p* values were calculated using Student’s *t*-test.

Upon identifying CmiCASP1, we therefore tested whether CmiCASP1 could cleave CmiGSDMA/B and trigger pyroptosis. Co-expression of CmiCASP1 and CmiGSDMA/B but not either protein alone in *GSDMD* and *GSDME* double-knockout HeLa cells (HeLa*^gsdmd/e DKO^* cells) led to pyroptotic morphology, featuring membrane blebbing and cell swelling and Annexin V/PI positivity in blebbing cells (**Figure 1G-H**). Moreover, co-expression CmiCASP1 and CmiGSDMA/B, but not either protein alone, in HEK293T cells also exhibited typical pyroptotic features, including increased Annexin V/PI double-positive cells (**Figure 1I**), enhanced LDH release (**Figure 1J**), and a marked reduction in intracellular ATP levels (**Figure 1K**). Western blot analysis showed that CmiCASP1-mediated cleavage of CmiGSDMA/B in HEK293T cells generated two N-terminal fragments (**Figure 1L**). Treatment with the pan-caspase inhibitor Z-VAD-FMK (Z-VAD) abrogated these pyroptotic morphology and CmiGSDMA/B cleavage (**Figure 1G-L**), confirming that this process is caspase-dependent.

### Cleavage of CmiGSDMA/B by CmiCASP1 occurred at D241 and D288

To identify the cleavage sites of CmiGSDMA/B by CmiCASP1, we analyzed the potential caspase-1 cleavage sites in CmiGSDMA/B based on the reported cleavage motifs of mammalian GSDMD and avian GSDMA, and finally identified D241 and D288 as potential sites (**Figure 2A**). We subsequently generated alanine substitution mutants at the two predicted caspase cleavage sites, including single mutants D241A and D288A, and double mutant D241A/D288A. As shown in **Figure 2B**, when co-transfected with CmiGSDMA/B into HEK293T cells, the D241A mutant abrogated production of the ∼30 kDa N-terminal fragment (p30, N241), whereas the D288A mutant eliminated the ∼35 kDa fragment (p35, N288). Meanwhile, the D241A/D288A double mutant completely blocked detectable cleavage of CmiGSDMA/B, indicating that D241 and D288 are the major cleavage sites of CmiCASP1 (**Figure 2B**). Notably, upon co-expression with CmiCASP1, HEK293T cells expressing D241A or D241A/D288A mutants but not the D288A mutant showed markedly attenuated pyroptosis compared to wild-type CmiGSDMA/B, as evidenced by reduced LDH release, a lower proportion of Annexin V^+^/PI^+^ cells and increased cell viability (**Figure 2C-E**). Consistently, fewer cells showing plasma membrane blebbing were observed when D241A or D241A/D288A mutants but not D288A mutant were co-expressed with CmiCASP1 in HeLa*^gsdmd/e DKO^* cells (**Figure 2F-G**). These findings demonstrate that the N241 fragment, generated by CmiCASP1-mediated cleavage of CmiGSDMA/B at D241, is essential for pyroptosis. Consistently, treatment with Z-VAD inhibited the cleavage of both D241A and D288A mutants into N288 fragment or N241 fragment, respectively (**Figure 2H**). Meanwhile, treatment with Z-VAD suppressed LDH release and the proportion of Annexin V^+^/PI^+^ cells, restored the cell viability, and inhibited membrane blebbing in cells expressing D288A mutant (**Figure 2I-L**). These findings reinforce the conclusion that both processing events of CmiGSDMA/B were dependent on CmiCASP1 and highlight D241 as a critical cleavage site for CmiGSDMA/B-mediated pyroptosis.

**Figure 2.**
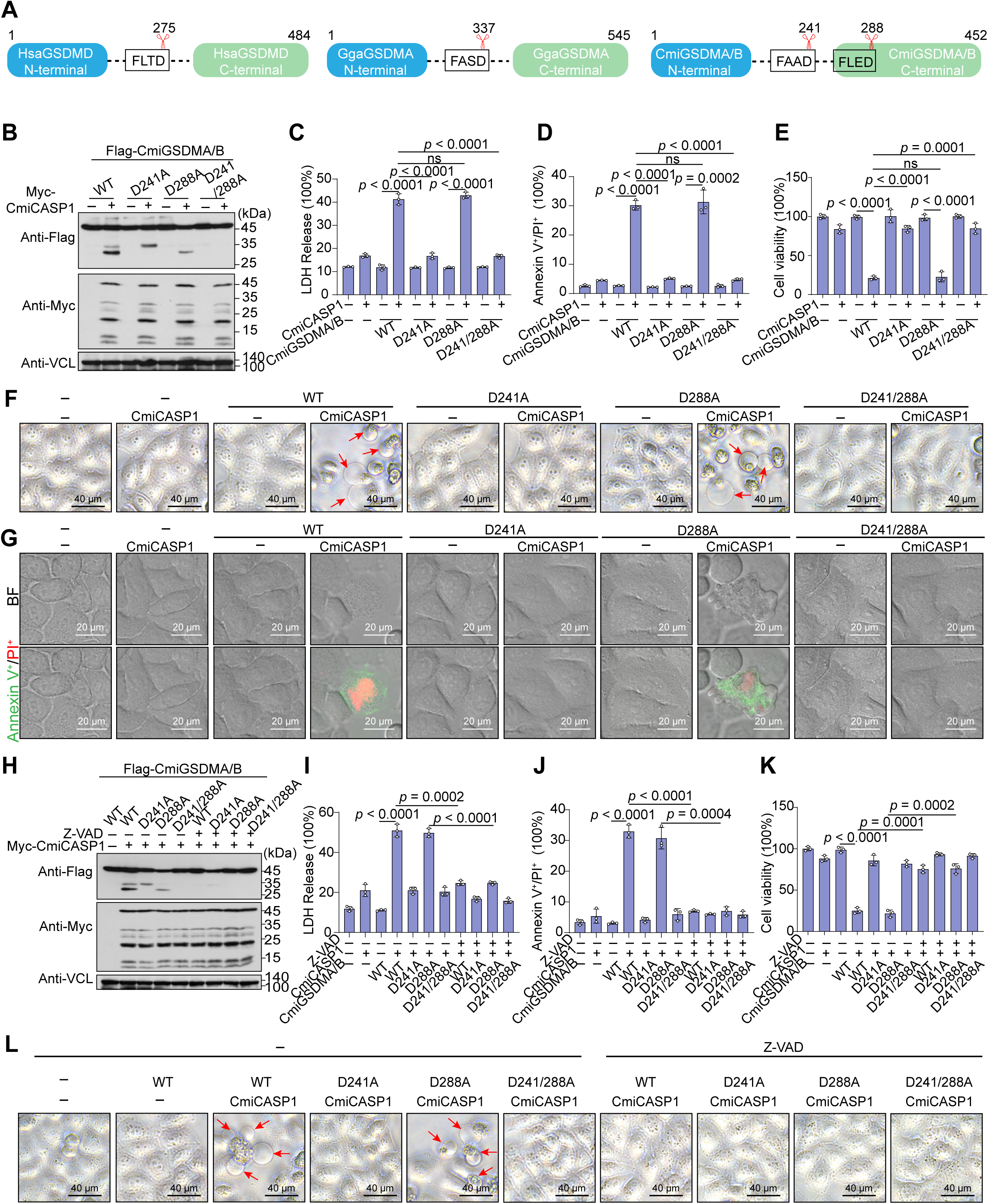
CmiGSDMA/B is cleaved by CmiCASP1 at D241 and D288. (**A**) Cartoon diagram showing the domain architecture of GSDMs and inflammatory caspase-mediated cleavage. **(B**) Western blot analysis of CmiCASP1-mediated cleavage of CmiGSDMA/B and its mutants in HEK293T cells. **(C)** LDH release assay in HEK293T cells co-expressing CmiCASP1 with CmiGSDMA/B or its mutants. **(D)** Flow cytometric analysis of Annexin V-FITC and PI double stained HEK293T cells transfected with the indicated constructs. (**E**) Cell viability assay of HEK293T cells transfected with the indicated expression vectors. (**F**) Bright-field images of HeLa*^gsdmd/e DKO^* cells co-transfected with CmiCASP1 and CmiGSDMA/B or its mutants. Red arrows indicate cells with pyroptotic morphology. (**G**) Microscopic analysis of cell death phenotypes by Annexin V-FITC/PI staining in HeLa*^gsdmd/e DKO^* cells under the indicated transfection conditions. **(H**) Western blot analysis of CmiCASP1-mediated cleavage of CmiGSDMA/B and its mutants in HEK293T cells treated with Z-VAD-FMK. **(I-K)** LDH release, flow cytometric analysis of Annexin V-FITC/PI staining, and cell viability assay of Z-VAD-FMK treated HEK293T cells transfected with the indicated expression vectors. (**L**) Bright-field images of HeLa*^gsdmd/e DKO^* cells under different transfection and treatment conditions, as indicated. Red arrows indicate cells with pyroptotic morphology. Data are shown as mean ± SD from three independent experiments. The *p* values were calculated using Student’s *t*-test.

### CmiGSDMA/B-N241 but not N288 can directly mediate pyroptosis

In line with D241 as a critical cleavage site for CmiGSDMA/B-mediated pyroptosis, expression of N241 but not N288 alone was sufficient to elicit pyroptosis in HeLa*^gsdmd/e DKO^* cells, as shown by cell swelling and plasma membrane blebbing (**Figure 3A-B**). Consistently, overexpression of N241 but not N288 alone in HEK293T cells resulted in increased Annexin V/PI double-positive cells, enhanced LDH release and decreased intracellular ATP levels (**Figure 3C-E**).

**Figure 3.**
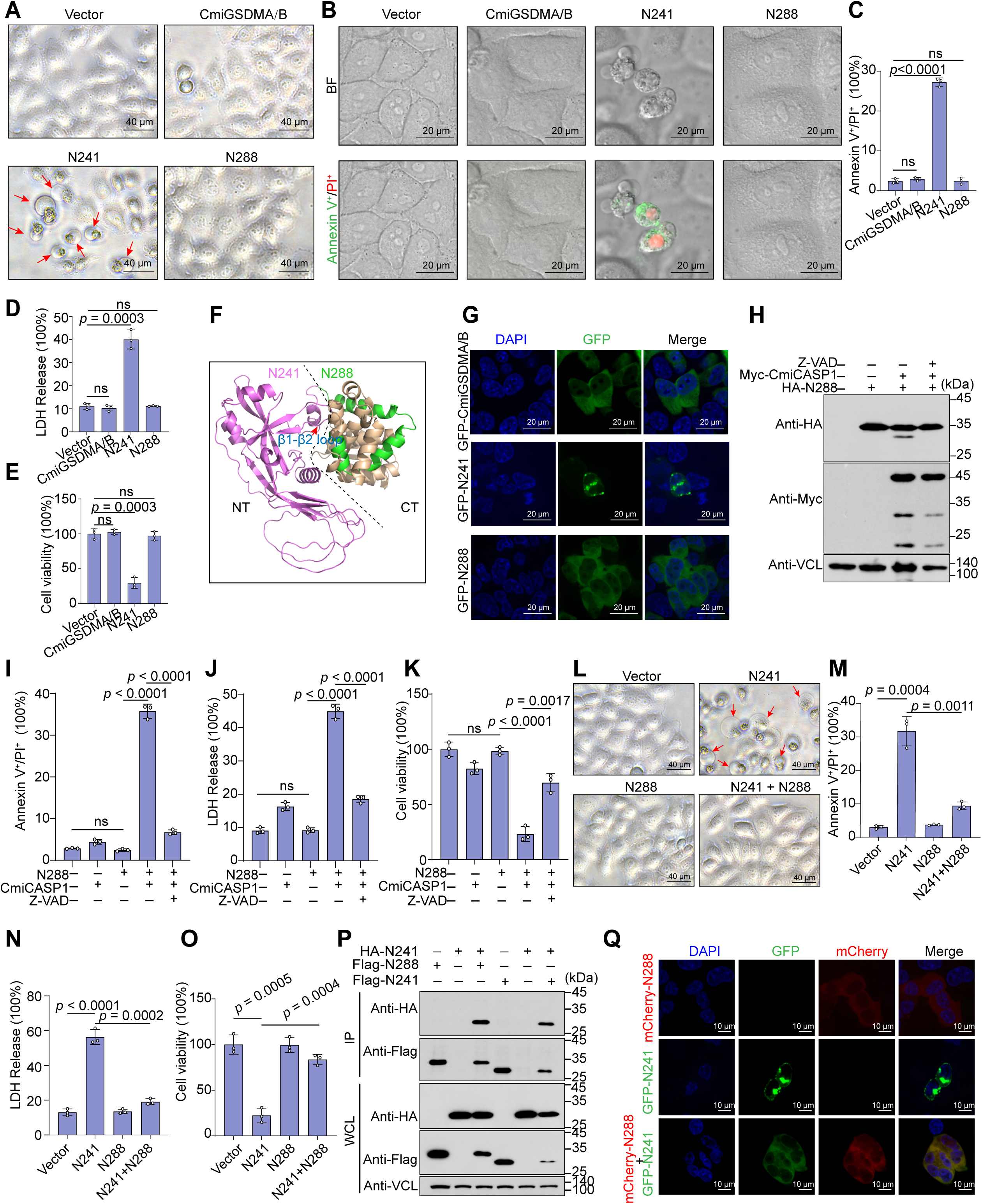
Distinct functional activities of N241 and N288. (**A**) Representative bright-field images of HeLa*^gsdmd/e DKO^* cells transfected with vectors expressing CmiGSDMA/B, N241 or N288. Red arrows indicate pyroptotic cells. (**B**) Microscopic analysis of cell death phenotypes by Annexin V-FITC/PI staining in HeLa*^gsdmd/e DKO^* cells transfected with expression vectors, as indicated. (**C-E**) Flow cytometry analysis of Annexin V-FITC and PI double-stained, LDH release assay, and ATP-based cell viability assay in HEK293T cells transfected with the indicated expression vectors. (**F**) The structure of CmiGSDMA/B predicted using AlphaFold. The N- and C-terminal domains are indicated and separated by a dashed line. The green region represents the additional segment present in N288 relative to N241 and the red triangle indicates the β1-β2 loop. (**G**) Subcellular localization of GFP-tagged N241 and N288 in HeLa*^gsdmd/e^ ^DKO^* cells. (**H**) Western blot analysis showing CmiCASP1-mediated cleavage of N288 in HEK293T cells and its inhibition by Z-VAD-FMK treatment. (**I-K**) Flow cytometry analysis of Annexin V-FITC/PI staining, LDH release assay, and cell viability assay in HEK293T cells co-expressing CmiCASP1 and N288 in the presence or absence of Z-VAD-FMK. (**L**) Representative bright-field images of HeLa*^gsdmd/e DKO^* cells expressing N241 alone or together with N288. Red arrows indicate cells with pyroptotic morphology. (**M-O**) Flow cytometry analysis of Annexin V-FITC/PI staining, LDH release assay, and cell viability assay in HEK293T cells expressing N241 alone or together with N288, as indicated. (**P**) Co-immunoprecipitation analysis of the interaction between N241 and N288 in HEK293T cells. (**Q**) Subcellular localization of GFP-tagged N241 in the presence or absence of mCherry-tagged N288 in HeLa*^gsdmd/e DKO^* cells. Quantitative data are presented as mean ± SD from three independent experiments. The *p* values were calculated using Student’s *t*-test.

To analyze the functional differences between the two fragments, we constructed a full-length structural model of CmiGSDMA/B using AlphaFold3. The model predicts that a single C-terminal extension of N288 (amino acids 242-288) folds back onto the N-terminal domain, yielding a compact self-inhibitory interface (**Figure 3F**). This region forms a molecular shroud masking the β1-β2 loop within the central domain of N241. In the cryo-EM structure of mammalian GSDMD, this β1-β2 loop was identified as an important membrane insertion motif for pore formation and pyroptosis (Xia *et al*., 2021). In line with the structural prediction, confocal microscopy showed that the N241 fragment localized to the plasma membrane (**Figure 3G**), similar to other gasdermin N-terminal domains. In contrast, the N288 fragment exhibited a diffuse cytoplasmic distribution without enrichment on the plasma membrane (**Figure 3G**). Since the N288 fragment contains a D241 cleavage motif, we further co-expressed the N288 fragment with CmiCASP1 in the presence or absence of Z-VAD. As shown, the N288 fragment could be further cleaved into N241, and such cleavage can be blocked by Z-VAD **(Figure 3H).** As expected, this cleavage further induced pyroptosis, as indicated by increased Annexin V/PI double-positive cells, massive LDH release, and decreased intracellular ATP levels (**Figure 3I-K**). Thus, N288 appears to represent an intrinsically inactive intermediate rather than a terminal effector fragment. Given that the additional 242-288 region was predicted to mask the β1-β2 loop and that N288 failed to enrich at the plasma membrane, we next tested whether N288 could suppress the activity of the active N241 fragment. Co-expression analysis showed that N288 could attenuate N241-mediated pyroptosis, as reflected by reduced Annexin V/PI double-positive cells, LDH release, and ATP depletion (**Figure 3L-O**). Co-immunoprecipitation (Co-IP) confirmed an interaction between N241 and N288 (**Figure 3P**), and fluorescence imaging showed that N288 may sequester N241, preventing its association with the cell membrane or its assembly into functional pores (**Figure 3Q**), suggesting a mechanism for N288-mediated inhibition of N241.

### LPS directly engages and activates CmiCASP1 via its CARD domain

After confirming that CmiGSDMA/B is a substrate of CmiCASP1, we investigated the upstream signals that activate CmiCASP1. In vertebrates, inflammatory caspases are typically engaged downstream of canonical or non-canonical inflammasomes (Ross *et al*., 2022; Zheng *et al*., 2020). To explore whether *C. milii* possesses similar upstream sensors, we searched its genome and proteome and identified 98 NLR (nucleotide-binding domain, leucine-rich repeat-containing) proteins, including two predicted proteins corresponding to a single NLRP3-like gene. Both proteins contain a fish-specific FISNA domain but lack an N-terminal PYD, and no CARD-containing NLR proteins were identified (**Figure 4—figure supplement 1A**). Additionally, no homologs of AIM2-like receptors or the ASC adaptor could be identified (**Figure 4—figure supplement 1A**). These findings indicated that *C. milii* lacks a complete recognizable canonical inflammasome signaling pathway, implying that CmiCASP1 may be activated by alternative upstream mechanisms.

**Figure 4.**
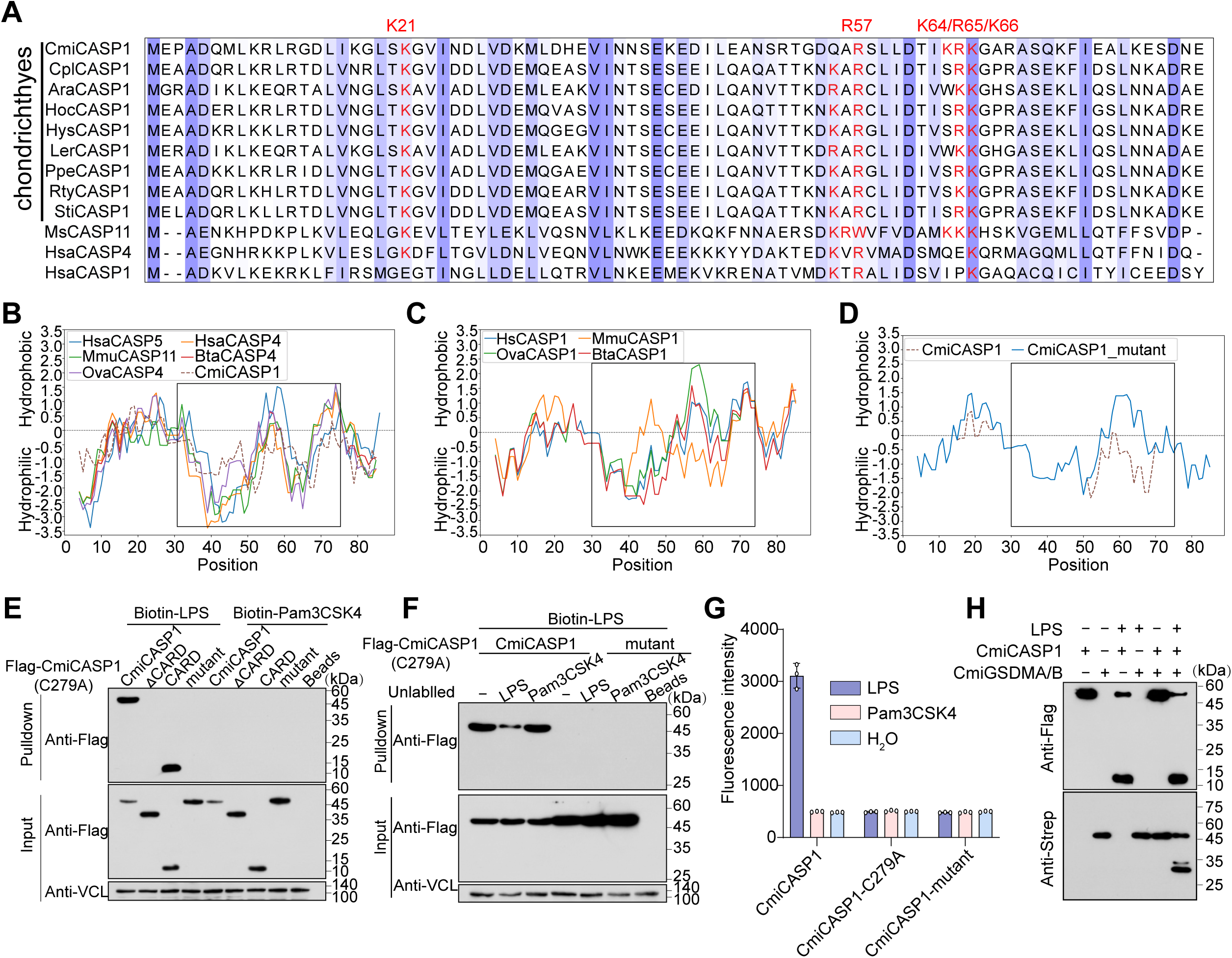
CmiCASP1 directly binds LPS to trigger activation. **(A)** Sequence alignment of caspase CARD domains generated using MAFFT. The conserved amino acid residues are highlighted in blue. Residues important for LPS binding to caspase-4/11 are highlighted in red. (**B-D**) Hydropathy plot of the CARD domains of mammalian caspase-1/4/5/11, CmiCASP1 and CmiCASP1-mutant generated using ProtScale. In the CmiCASP1-mutant, conserved positively charged residues within the predicted LPS-binding surface were replaced by alanine. Dashed box indicates the distinctive hydrophobic grooves within CARD domains of caspase-1 and caspase-4 homologs and CmiCASP1. (**E**) Streptavidin-based pull-down assay to detect biotin-conjugated LPS binding to Flag-tagged catalytically dead CmiCASP1-C279A, CmiCASP1-C279A-ΔCARD, the isolated CARD domain, and the CmiCASP1-C279A-mutant in transfected HEK293T cell lysates. (**F**) Streptavidin-based pull-down assay to detect the binding of biotin-labeled LPS or Pam3CSK4 to Flag-tagged CmiCASP1-C279A and CmiCASP1-C279A-mutant, with the indicated unlabeled ligands for competition experiments. (**G**) LPS induced activation of purified CmiCASP1. Caspase activity was determined by measuring the fluorescence intensity of free AMC hydrolyzed from Z-VAD-AMC. (**H**) Purified CmiGSDMA/B was cleaved in vitro by CmiCASP1, which was activated by LPS.

Besides the canonical inflammasome signaling pathway, mammals possess a non-canonical inflammasome pathway mediated by caspase-4/5/11. The non-canonical pathway enables direct sensing of cytosolic lipopolysaccharide (LPS) through the CARD domains of caspase-4/5/11, as mutation analyses demonstrate that substituting specific positively charged residues abolishes LPS binding (Shi *et al*., 2014). Sequence alignments revealed that the CARD domain of chondrichthyan CASP1 shares conserved LPS-binding positively charged residues with mammalian caspase-4/11 (**Figure 4A**). In addition, hydrophobicity analysis identified two hydrophobic regions reminiscent of the two LPS-binding hydrophobic grooves described in the HyCaspA study (**Figure 4B-D**) (Chen *et al*., 2023). These observations prompted us to test the possibility of a non-canonical inflammasome-like activation mechanism in *C. milii*.

To avoid potential interference from caspase auto-processing during LPS-binding assays, we constructed the catalytically inactive CmiCASP1-C279A mutant. Based on the CmiCASP1-C279A mutant, we further generated a CARD-deletion mutant (ΔCARD) and a multisite mutant, in which conserved positively charged residues within the predicted CARD LPS-binding surface were replaced by alanine (CmiCASP1-C279A-mutant). Biotinylated LPS efficiently precipitated CmiCASP1-C279A, whereas biotinylated Pam3CSK4 did not (**Figure 4E**). Specificity was confirmed by competition experiments with excess unlabeled LPS, which significantly reduced CmiCASP1 binding but had no effect on Pam3CSK4 (**Figure 4F**). Importantly, deletion of the CARD domain (ΔCARD) or mutation of the conserved positively charged residues in the C279A background (CmiCASP1-C279A-mutant) abolished detectable binding to LPS (**Figure 4E**), suggesting that the CARD domain is essential for CmiCASP1 to recognize LPS.

Having established CARD-dependent LPS binding by CmiCASP1, we next examined whether this interaction was sufficient to enhance its protease activity and promote downstream substrate processing. Using the fluorogenic substrate Z-VAD-AMC, we found that CmiCASP1 exhibited only weak basal activity, but incubation with LPS significantly increased its protease activity consistent with CmiCASP1 activation (**Figure 4G**). In contrast, the CmiCASP1-C279A mutant and CmiCASP1-mutant did not respond to LPS (**Figure 4G**). Moreover, to assess the function of LPS-mediated CmiCASP1 activation, we incubated LPS-stimulated CmiCASP1 with CmiGSDMA/B.

Upon LPS treatment, CmiCASP1 efficiently cleaved CmiGSDMA/B, generating two distinct N-terminal fragments (**Figure 4H**), corresponding to the generation of the previously identified cleavage products. Together, these results indicate that LPS binding through the CARD domain enhances CmiCASP1 activity and promotes CmiGSDMA/B processing.

### N241 of CmiGSDMA/B exhibits bactericidal activity

Previous studies have shown that gasdermins can contribute to antibacterial defense by directly damaging bacterial membranes. In particular, GSDMD has been reported to exert bactericidal activity against bacteria (Liu *et al*., 2016), whereas the antibacterial activity of human GSDMB remains controversial and appears to depend on isoform composition, proteolytic processing and experimental context (Hansen *et al*., 2021; Wang *et al*., 2019). We therefore asked whether antibacterial activity is retained in cartilaginous fish GSDMA/B. To this end, full-length CmiGSDMA/B and its fragments N241 and N288 were expressed in HEK293T cells. Cell lysates were harvested, concentrated, and incubated with different bacterial strains. Bacterial viability was determined by colony formation assay (CFU assay). The lysate containing N241 significantly inhibited colony formation of Gram-negative bacteria, including *Escherichia coli*, *Edwardsiella tarda* strain EIB202, and *Vibrio parahaemolyticus* but had no inhibitory effect on Gram-positive bacteria such as *Staphylococcus aureus*, *Listeria monocytogenes*, and *Bacillus megaterium* (**Figure 5A-B** and **Figure 5—figure supplement 1A**). Under the same conditions, lysates containing full-length CmiGSDMA/B or N288 did not show antimicrobial activity (**Figure 5A-B** and **Figure 5—figure supplement 1A**).

**Figure 5.**
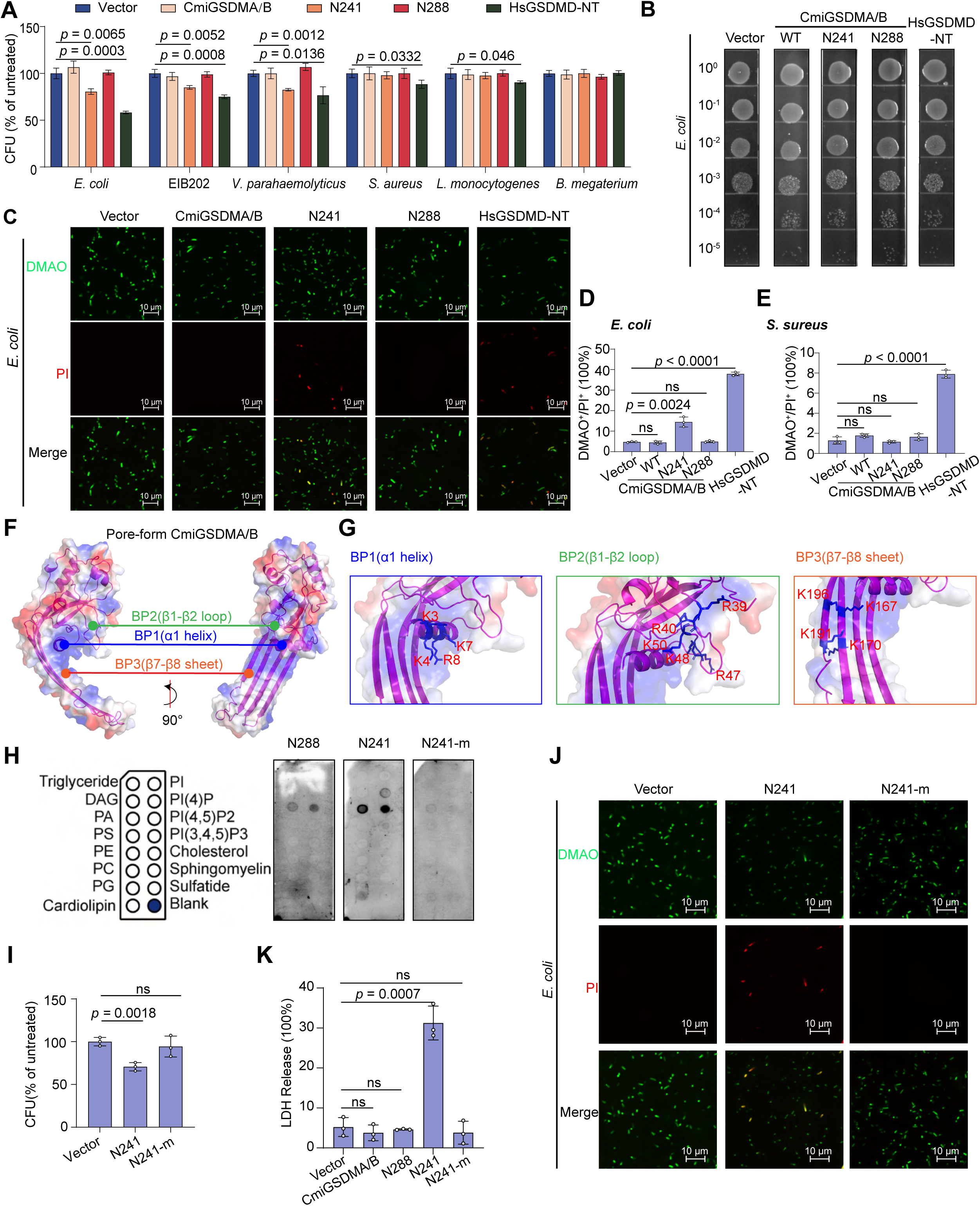
N241 exhibits antibacterial activity against Gram-negative bacteria. (**A**) Effect of CmiGSDMA/B and its fragments on bacterial viability. HEK293T cell lysates containing full-length CmiGSDMA/B, N241 or N288 fragments were incubated with Gram-negative bacteria (*E. coli*, *E. tarda* strain EIB202, and *V. parahaemolyticus*) and Gram-positive bacteria (*S. aureus*, *L. monocytogenes*, and *B. megaterium*). Bacterial viability was assessed by CFU assays. Only plates containing 30-300 colonies were used for CFU calculation. (**B**) Representative serial dilution plating of *E. coli* treated with the indicated GSDM-containing lysates. *E. coli* suspended in PBS was incubated with the indicated lysates at 37°C for 40 minutes, serially diluted 10-fold and plated on LB agar for CFU determination. (**C**) Representative fluorescence microscopy images showing the viability of *E. coli* assessed by DMAO/PI staining. Bacteria were incubated for 40 minutes with the indicated concentrated lysates collected from transfected HEK293T cells. DMAO permeates both live and dead bacteria, while PI stains only dead bacteria. (**D, E**) Flow cytometry analysis of DMAO and PI double-stained *E. coli* (**D**) or *S. aureus* (**E**) after incubation with the indicated concentrated lysates. (**F, G**) Structural analysis of N241 highlights putative membrane contact and insertion surfaces enriched in basic residues including three basic patches (BPs). (**H**) Lipid strip binding assay compared the differences in lipid binding between purified wild-type N241, N288, and N241-m. (**I**) CFU assay measuring the viability of *E. coli* after incubation with lysates containing N241 and N241-m. (**J**) Representative fluorescence microscopy images showing the viability of *S. aureus* assessed by DMAO/PI staining. (**K**) LDH release assay of HEK293T cells transfected with the indicated vectors. Quantitative data are presented as mean ± SD from three independent experiments. The *p* values were calculated using Student’s *t*-test.

To distinguish between simple inhibitory effects and bactericidal effects, the viability of bacteria was determined by investigating the integrity of the cell membrane using dual-fluorescence staining with DMAO and PI. *E. coli* and *S. aureus* were chosen as representative Gram-negative and Gram-positive species. The results showed that lysates containing the N241 fragment significantly increased the proportion of PI-positive *E. coli* compared with control lysates, indicating damage to bacterial cell membrane, whereas full-length CmiGSDMA/B and N288 had no effect (**Figure 5C**). Flow cytometry confirmed that about 15% of *E. coli* treated with N241-containing lysates were PI positive (**Figure 5D**). For *S. aureus*, however, lysates containing the N241 fragment showed no significant PI uptake compared with vector, full-length CmiGSDMA/B or N288 lysates (**Figure 5E** and **Figure 5—figure supplement 1B**). These results suggest that N241 exhibits specific bactericidal activity against Gram-negative bacteria.

Since the surfaces of Gram-negative bacteria are rich in negatively charged lipids, and electrostatic attraction is a common mechanism for antimicrobial peptides to bind bacterial membranes (Brogden, 2005), we next investigated whether the basic residues in N241 involved in membrane insertion are required for bactericidal activity. Upon structural modeling, we identified three basic regions potentially involved in membrane integration, the α1 helix (base region 1, BP1), the β1-β2 loop (base region 2, BP2), and the β7-β8 loop structure (base region 3, BP3) (**Figure 5F-G**). We then generated an N241 mutant, termed N241-m, in which Lys/Arg residues were substituted with Ala (**Figure 5—figure supplement 1C**). Lipid-binding assays showed that wild-type N241 preferentially bound PtdIns(4)P, PtdIns(4,5)P2, phosphatidic acid (PA), and cardiolipin (**Figure 5H**). Binding to PtdIns(4,5)P2 and PA is consistent with previous observations in other GSDM family members and supports the hypothesis that N241 targets cytoplasmic leaflets of membranes (Balla, 2013; Liu *et al*., 2016). Cardiolipin is a characteristic lipid of mitochondrial inner membranes and is also found in bacterial membranes (Li *et al*., 2015). The binding of N241 to cardiolipin provides a plausible explanation for its interaction with Gram-negative bacteria. In contrast, N241-m showed significantly reduced binding to these acidic phospholipids (**Figure 5H**). Functional assays confirmed that N241-m lost bactericidal activity against *E. coli*, and failed to induce pyroptosis (**Figure 5I-K**). These results indicate that the positively charged regions of N241 are critical for its membrane-targeting and are required for N241-mediated pyroptosis and bactericidal activity.

## Discussion

In mammals, cytosolic LPS triggers the non-canonical inflammasome pathway through direct sensing by inflammatory caspase-4/5/11 (Shi *et al*., 2014), which can cleave GSDMD to trigger pyroptosis (Kayagaki *et al*., 2015; Shi *et al*., 2015). In this study, we focused on cartilaginous fish *C. milii*, a representative of the ancestral jawed vertebrate lineage (Venkatesh *et al*., 2014), whose CmiGSDMA/B occupies a basal position within the vertebrate GSDMA/B/C/D family (Wang *et al*., 2023b). We found that CARD-dependent LPS recognition activates CmiCASP1, which cleaves CmiGSDMA/B to generate an N241 fragment that induces pyroptosis, establishing a functional connection between bacterial LPS sensing, inflammatory caspase activation and a GSDMA/B/C/D subfamily effector in the early jawed vertebrates (**Figure 6**). In parallel, canonical inflammasomes generally comprise a cytosolic sensor, ASC and pro-caspase-1, leading to caspase-1 activation and maturation of pro-IL-1β and pro-IL-18 (Fu *et al*., 2024; Rathinam and Fitzgerald, 2016). Importantly, cartilaginous fishes also represent an early vertebrate lineage in which recognizable pro-IL-1β and pro-IL-18 can be detected (**Figure 4—figure supplement 1B-C**) (Bird *et al*., 2002; Rivers-Auty *et al*., 2018). Notably, whereas *C. milii* appears to lack the key molecular machinery for canonical inflammasomes (**Figure 4—figure supplement 1A**), other cartilaginous fish lineages, exemplified by *Chiloscyllium plagiosum,* appeared to contain these key components, such as NLRC4 and ASC (**Figure 4—figure supplement 1D**). Moreover, the genes encoding these components show broadly similar expression patterns (**Figure 4—figure supplement 1E**). In addition, *C. plagiosum* pro-IL-1β and pro-IL-18 contain predicted caspase-processing sites (**Figure 4—figure supplement 1B-C**). These comparisons further implied that at the early jawed vertebrates, both non-canonical and canonical inflammatory pathways have been established.

**Figure 6.**
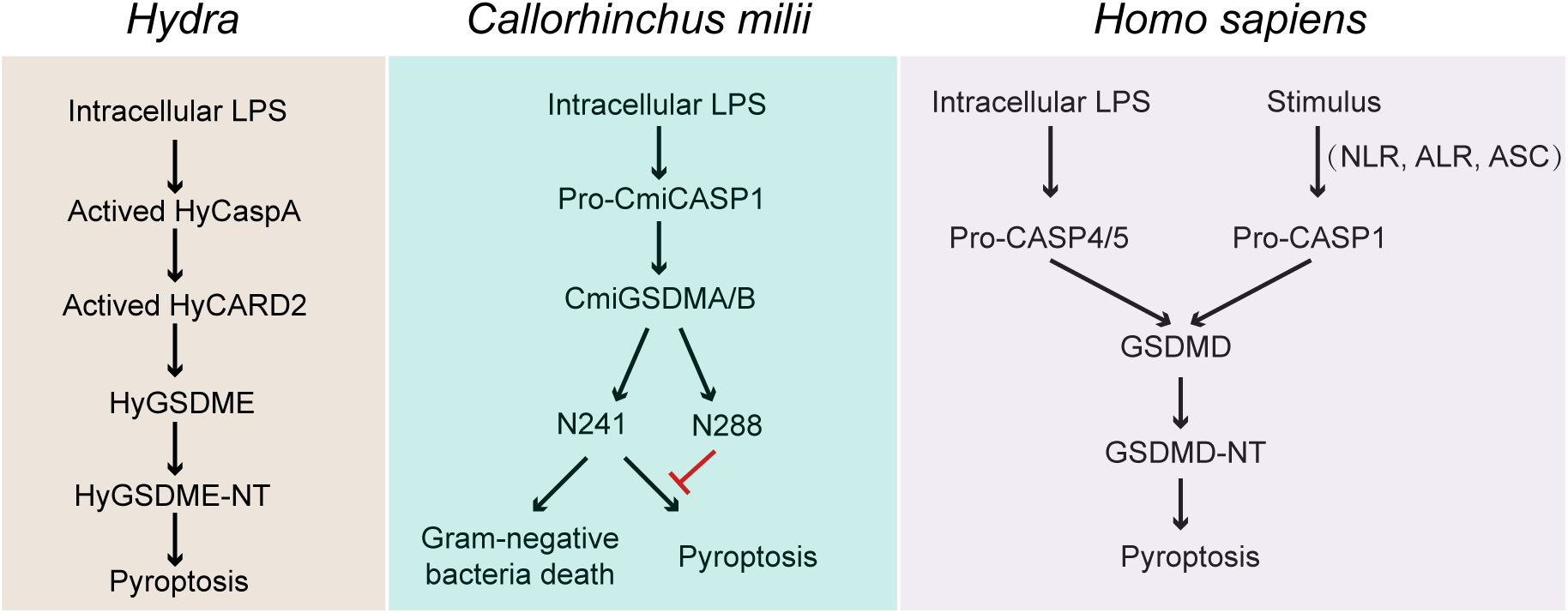
Working model of LPS-responsive CmiCASP1-CmiGSDMA/B pathway. Schematic model of the CmiCASP1-mediated non-canonical pyroptosis pathway in *C. milii*, highlighting its potential contributions to antibacterial immune responses.

Besides vertebrate GSDMA/B/C/D clade, an LPS-responsive caspase-gasdermin pathway has also been reported in *Hydra*, where cytosolic LPS activates a caspase-dependent pyroptotic response mediated by GSDME (Chen *et al*., 2023). Previous studies have suggested that, within the vertebrate GSDMA/B/C/D lineage, caspase-1-dependent pyroptosis is mainly mediated by GSDMA in non-mammalian tetrapods (Billman *et al*., 2024), whereas this role is predominantly associated with GSDMD in mammals (Kayagaki *et al*., 2015; Shi *et al*., 2015). Our finding that CmiCASP1 cleaves CmiGSDMA/B places a GSDMA/B/C/D effector downstream of an inflammatory caspase in a cartilaginous fish lineage. This configuration raises the possibility that inflammatory caspases were coupled to GSDMA/B or GSDMA effectors before the predominant association with GSDMD in mammals. These comparisons reflect the convergent evolution in LPS-triggered pyroptosis, wherein distinct gasdermin family members are deployed as executioners across different phyla (**Figure 6**).

In addition to convergent evolution in upstream activator and downstream gasdermin substrate (Billman *et al*., 2024; Chen *et al*., 2021; Xue *et al*., 2019), CmiGSDMA/B reveals a regulatory mode in which a single inflammatory caspase generates cleavage products with divergent activities. CmiCASP1-mediated cleavage of CmiGSDMA/B generates two N-terminal fragments with different activities, among which N241 acted as the active fragment that induced pyroptosis, whereas N288 failed to mediate pyroptosis and attenuated N241-mediated cell death. Such mode of regulation is reminiscent of amphioxus GSDME (Wang *et al*., 2023b). Additionally, in corals and mollusks, cleavage of GSDME by caspase-3 also produces two N-terminal fragments with similar pyroptotic activity (Jiang *et al*., 2020; Qin *et al*., 2023). Therefore, multi-site cleavage is probably an ancient and flexible regulatory mechanism found in gasdermin lineages. CmiGSDMA/B represents an early vertebrate example in which a single inflammatory caspase generates both an active and a regulatory fragment, thereby fine-tuning gasdermin activity prior to the specialization of vertebrate gasdermin paralogs.

Previous studies have shown that mammalian GSDMD-NT can directly damage bacterial membranes and kill Gram-negative bacteria as well as some Gram-positive bacteria (Liu *et al*., 2016; Wang *et al*., 2019), whereas human GSDMB-NT has been reported to kill selected Gram-negative bacteria (Hansen *et al*., 2021; Wang *et al*., 2023a). In this study, N241 fragment disrupted the membrane integrity of *E. coli* but was ineffective against *S. aureus*. This selective activity may be related to differences in the lipid composition of bacterial membrane, the type of surface charge or the physical barrier imposed by the thick peptidoglycan layer of Gram-positive bacteria (Li *et al*., 2017; Silhavy *et al*., 2010; Sohlenkamp and Geiger, 2016). Since the positively charged region of N241 is required for binding to acidic lipids, pyroptotic activity and antibacterial activity, the N241-mediated pyroptosis and antibacterial activity may share the same membrane-targeting strategy (Liu *et al*., 2016). Thus, dual outputs may be particularly relevant in an early vertebrate context, in which inflammatory caspase activation could couple host-cell membrane disruption with direct damage to susceptible bacterial membranes.

## Materials and Methods

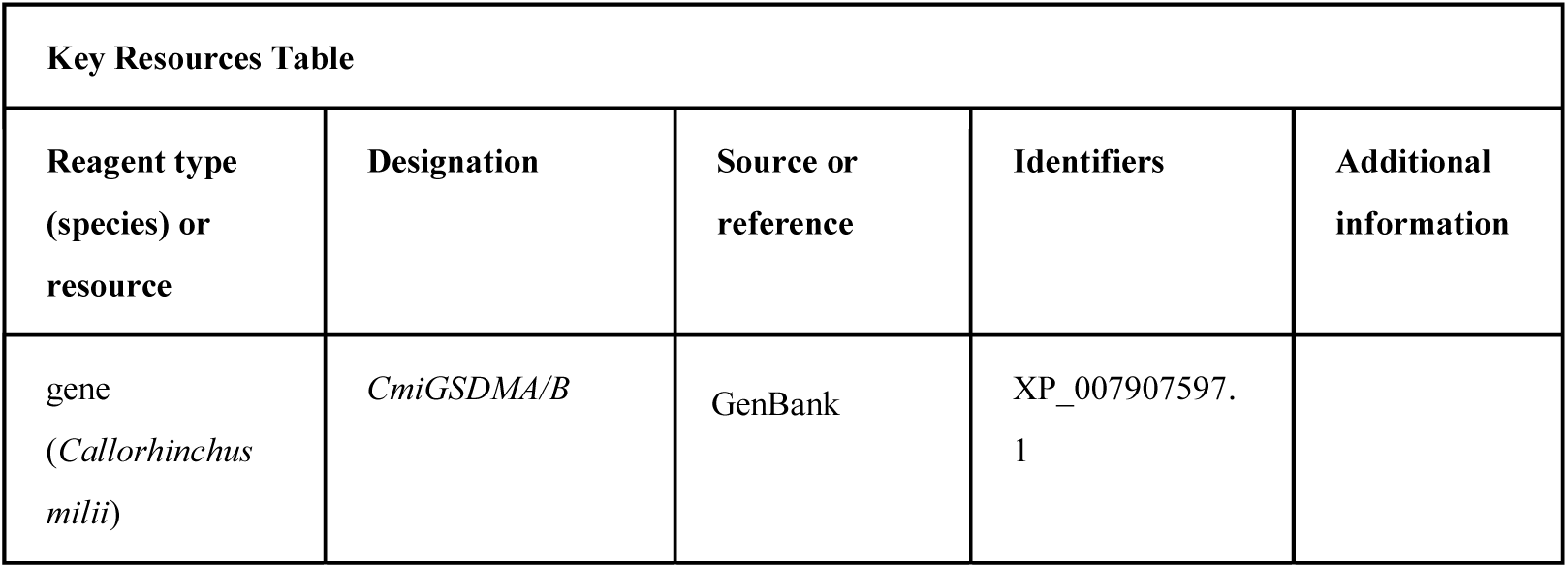
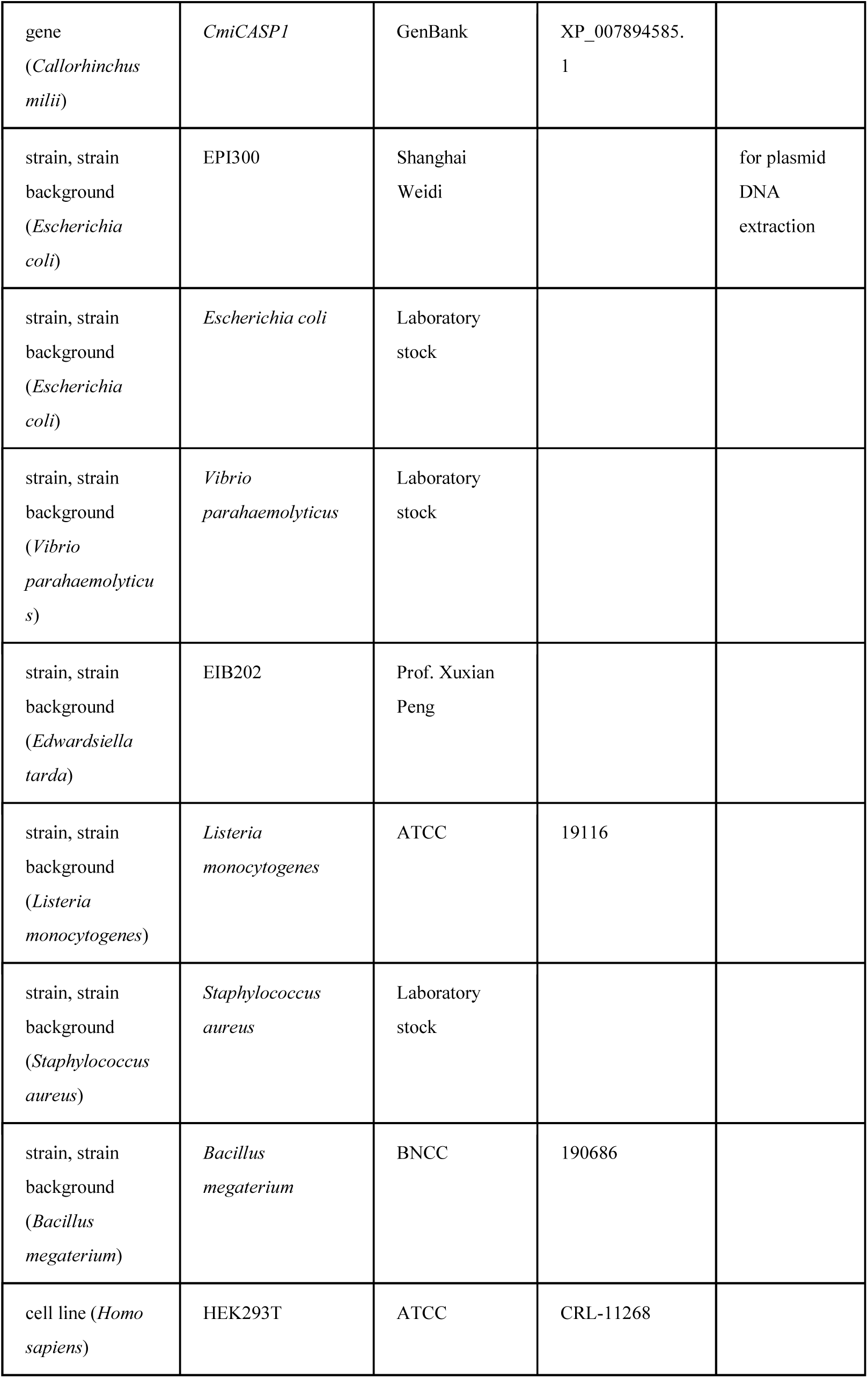
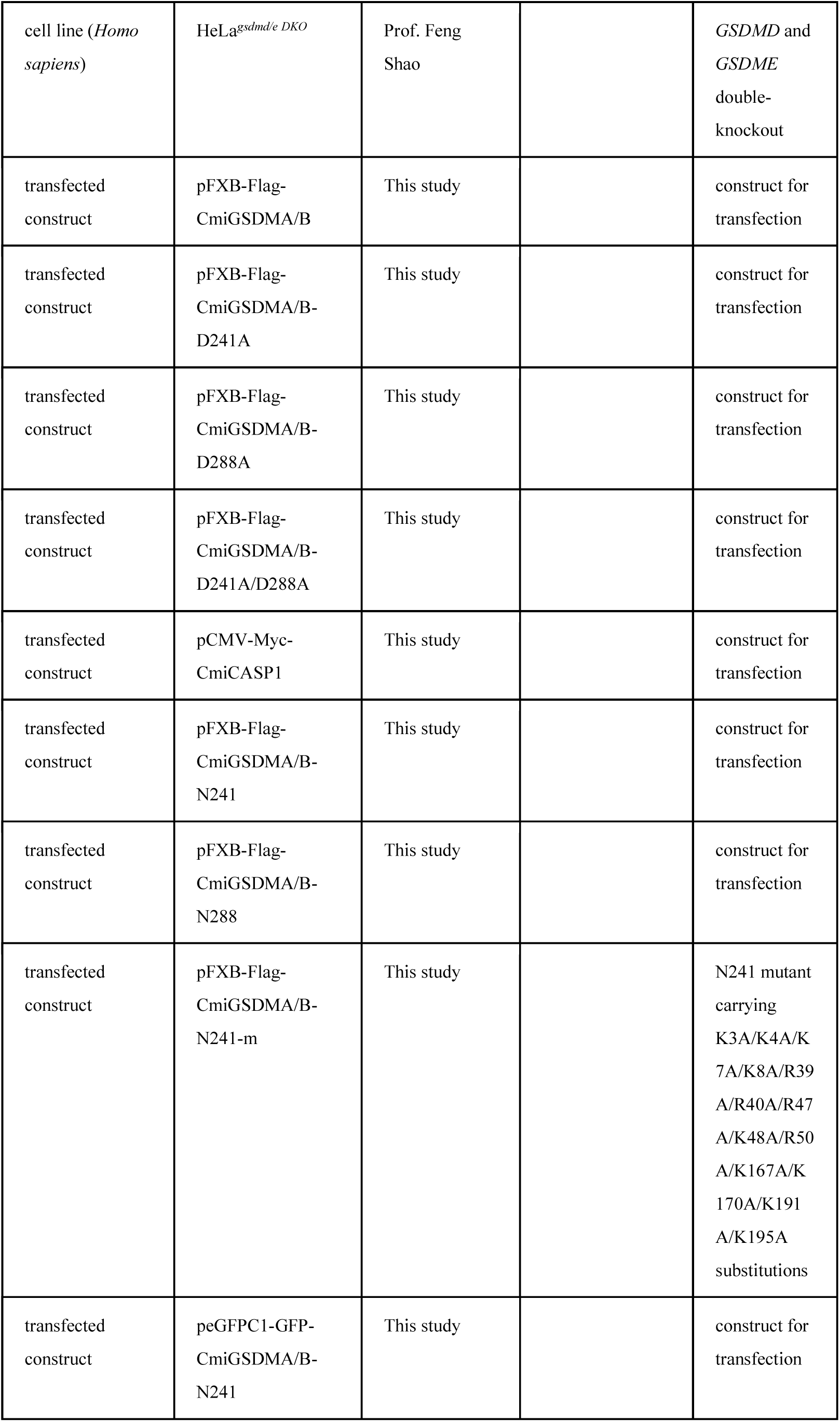
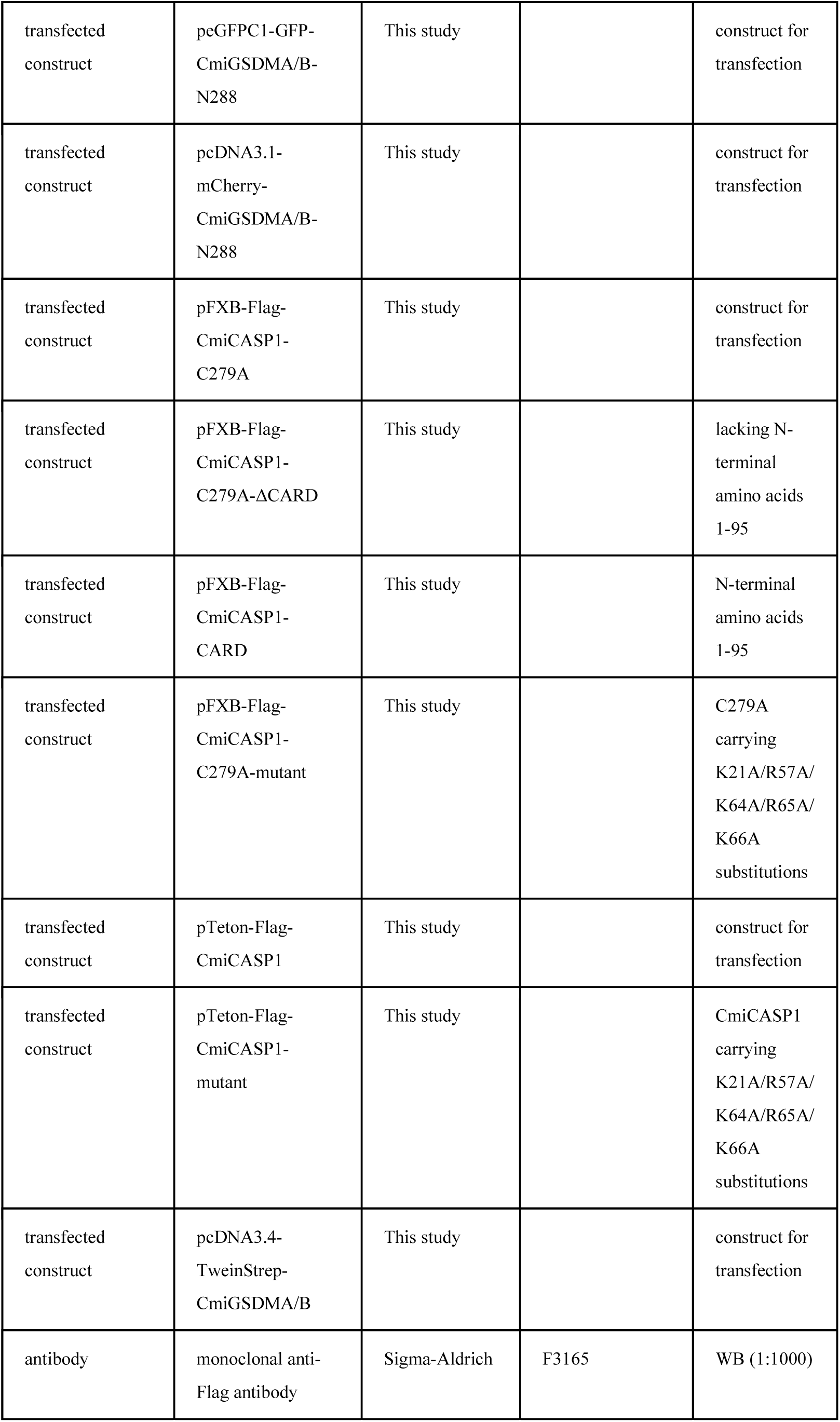
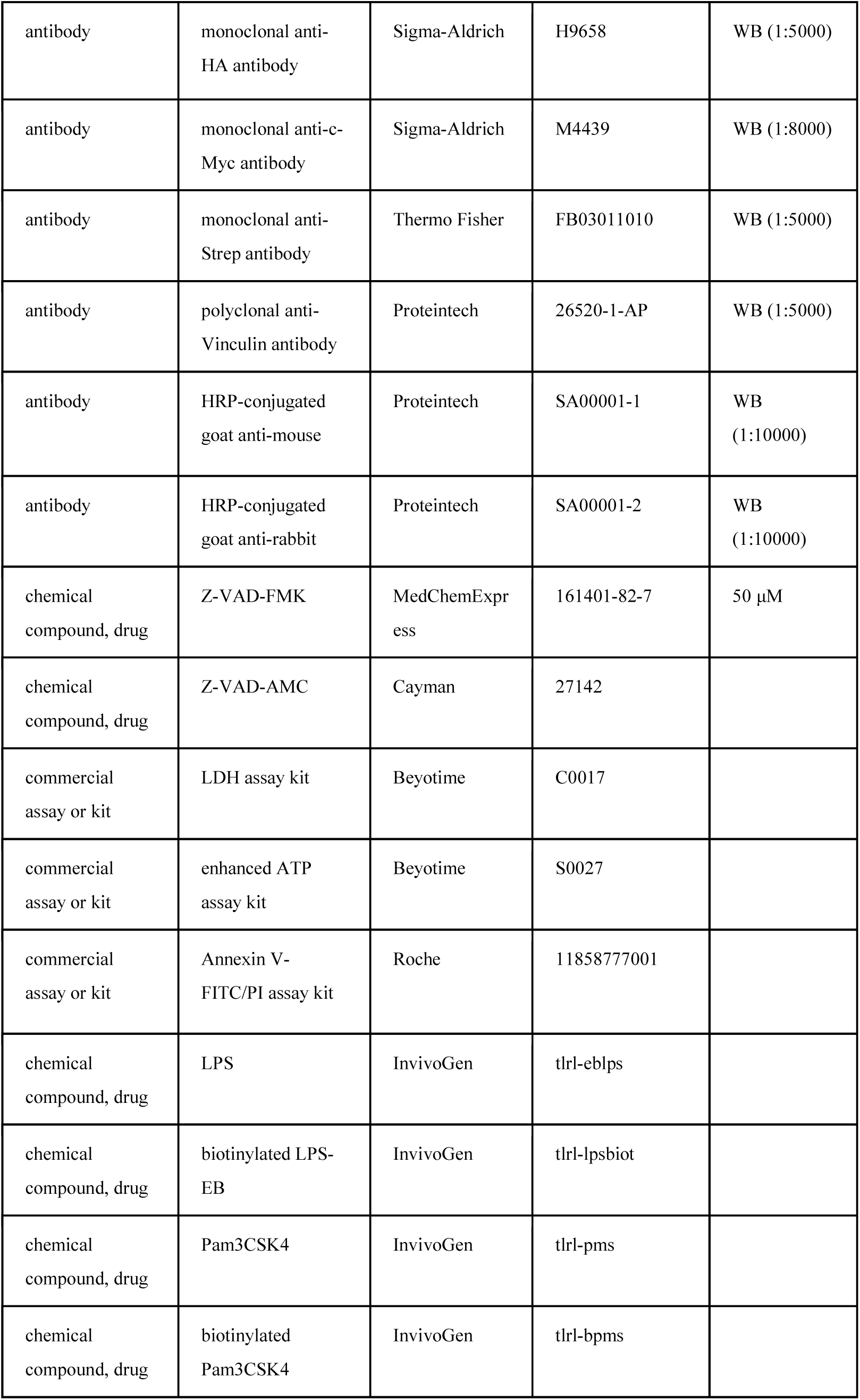
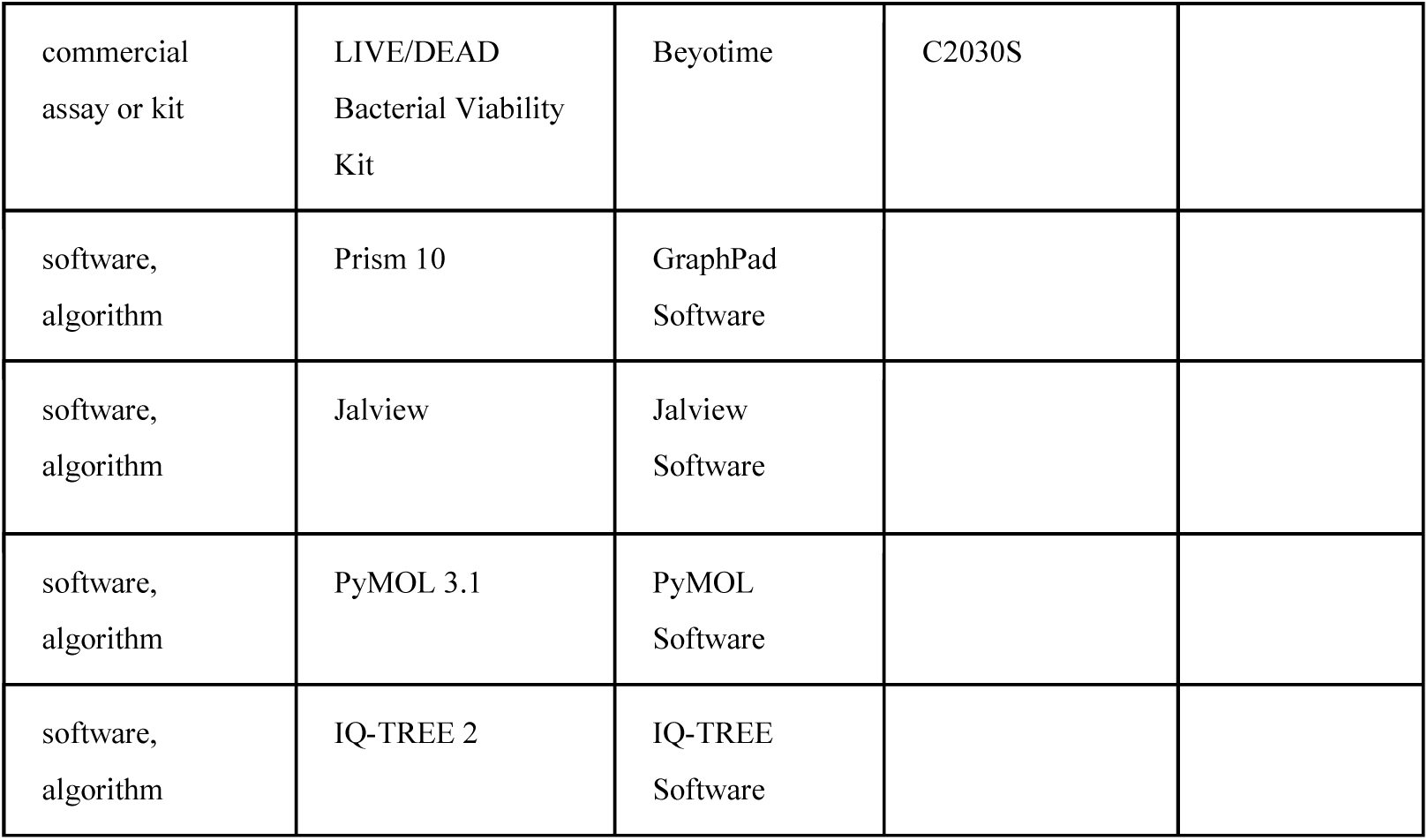

### Cell lines and bacterial strains

HEK293T were obtained from the American Type Culture Collection (ATCC). HeLa*^gsdmd/e DKO^* cells were kindly offered by Prof. Feng Shao (National Institute of Biological Sciences, Beijing, China) (Wang *et al*., 2017). These cells were cultured in Dulbecco’s modified Eagle Medium (DMEM, Gibco), supplemented with 10% fetal bovine serum (FBS, Gibco), at 37°C in a 5% CO_2_ humidified incubator. Both cell lines were tested negative for mycoplasma, and HEK293T were authenticated by STR profiling. *Escherichia coli* (*E. coli*) was grown in Luria Broth (LB). *Staphylococcus aureus* (*S. aureus*) and *Edwardsiella tarda* strain EIB202 were cultured in Tryptic Soy Broth (TSB). *Listeria monocytogenes* (*L. monocytogenes*) was cultured in Brain-Heart Infusion (BHI) medium, and *Bacillus megaterium* (*B. megaterium*) was maintained in Nutrient Broth (NB). *Vibrio parahaemolyticus* (*V. parahaemolyticus*) was grown in Marine Luria Broth (MLB).

### Sequence and phylogenetic analysis

Protein sequences were retrieved from the National Center for Biotechnology Information database. Multiple sequence alignments were performed using MAFFT (Katoh *et al*., 2002), and visualized with Jalview (Procter *et al*., 2021). For phylogenetic analysis, alignments of representative protein sequences were trimmed to remove poorly aligned regions. Maximum-likelihood phylogenetic trees were inferred using IQ-TREE with ModelFinder to select the best-fit amino acid substitution model. Branch support was evaluated using 5,000 ultrafast bootstrap replicates (Nguyen *et al*., 2015). The resulting phylogenetic trees were visualized using FigTree and further formatted for presentation.

### Plasmid constructions and transfection

The protein coding sequences (CDSs) of *CmiGSDMA/B* (GenBank accession no. XP_007907597.1) and *CmiCASP1* (GenBank accession no. XP_007894585.1) were chemically synthesized by Tsingke Biotech (Beijing, China) and subcloned into a cloning vector. The CDSs of CmiGSDMA/B-N241 (1 to 241 residues), CmiGSDMA/B-N288 (1 to 288 residues) were subcloned on the basis of the synthesized sequences. Site-directed mutagenesis of CmiGSDMA/B Asp241 to Ala (D241A), Asp288 to Ala (D288A) and the double-sites mutant of D241A/D288A were performed using the QuikChange Lightning kit (Cat. 210503, Agilent) according to the manufacturer’s instructions. CmiCASP1 C279A and CmiCASP1 multi-site mutants were generated using the same kit. For transfection, the CDSs were subcloned into Flag-tagged, Myc-tagged or Strep-tagged expression vectors. JetPRIME transfection reagent was used for transient transfection of plasmids into cells.

### Protein purification

For Flag-tagged protein purification, cells expressing target gene were collected and washed twice by ice-cold PBS. Cells were lysed in the buffer A containing 20 mM Tris (pH 7.5), 150 mM NaCl, 1 mM EDTA, 0.1% Triton X-100 (v/v), and protease inhibitor cocktail (Cat. 4693132001, Roche) for 30 min. Cell lysates were clarified by centrifugation at 12,000 × *g* for 10 min at 4°C and the supernatants were incubated with anti-Flag resins (Cat. A2220, Sigma-Aldrich) overnight at 4℃. The resins were washed five times with lysis buffer and three times with buffer B (20 mM Tris (pH 7.5), 150 mM NaCl, 1 mM PMSF). The Flag-tagged proteins were then eluted using 3 × Flag peptides (Cat. F4799, Sigma-Aldrich) according to the manufacturer’s protocol. The purified proteins were analyzed by SDS-PAGE and Coomassie brilliant blue staining to assess purity.

For Strep-tagged proteins, cells expressing N-terminally Strep-tagged proteins were collected and washed twice with ice-cold PBS. Cells were resuspended in lysis buffer containing 100 mM Tris (pH 7.5), 150 mM NaCl, 1 mM EDTA and protease inhibitor cocktail, and then disrupted by sonication. To remove free biotin, avidin was added to the lysates, followed by incubation on ice for 30 min. Cell lysates were clarified by centrifugation at 18,000 × *g* for 5 min at 4°C. The supernatants were loaded onto a Strep-Tactin column. After gravity-flow binding, the column was washed with five column volumes of lysis buffer. Bound proteins were eluted with buffer E (100 mM Tris (pH 8.0), 150 mM NaCl, 1 mM EDTA, 2.5 mM desthiobiotin) according to the manufacturer’s protocol. The purified proteins were analyzed by SDS-PAGE and Coomassie staining to assess purity.

### Microscopy imaging

To examine cell death morphology, HeLa*^gsdmd/e DKO^* cells were plated in 12-well plates overnight, transfected with the indicated plasmids using jetPRIME transfection reagent. Bright-field cell images were captured using a microscope (Leica Microsystems CMS GmbH DMi8). To assess plasma membrane permeability and cell death, cells transfected with the indicated plasmids were stained using the Annexin V-FITC/PI Apoptosis Assay Kit according to the manufacturer’s instructions. To examine the subcellular localization of N241 and N288, HeLa*^gsdmd/e^ ^DKO^* cells were transfected with the indicated GFP-tagged or mCherry-tagged plasmids. Fluorescence images were acquired using a Nikon Eclipse Ti2 confocal microscope.

### Flow cytometry

HEK293T cells (1.5 × 10^5^ cells per well) were seeded in 24-well plates and cultured overnight. Cells were transfected with the indicated plasmids and incubated in fresh Opti-MEM (Cat. 31985070, Thermo Fisher) for 20 h as described above. After incubation, cells were collected and washed with ice-cold PBS, then double-stained with Annexin V-FITC and PI according to the manufacturer’s instructions, and analyzed by flow cytometry (Beckman Coulter Cyto-FLEX S).

### Cell viability and cytotoxicity assays

HEK293T cells (1.5 × 10^5^ cells per well) were seeded in 24-well plates and cultured overnight. Cells were transfected with the indicated plasmids, followed by incubation in fresh Opti-MEM for 20 h. Intracellular ATP levels were measured using the Enhanced ATP Assay Kit according to the manufacturer’s instructions and used as an indicator of cell viability.

For LDH release assays, HEK293T cells (4 × 10^4^ cells per well) were seeded in 48-well plates and cultured. After transfection with the indicated plasmids, cells were incubated in fresh Opti-MEM for 20 h. Culture supernatants were collected, and LDH release was measured using the LDH Release Assay Kit according to the manufacturer’s instructions. Cytotoxicity was calculated using the following equation: cytotoxicity (%) = 100 × (experimental sample − medium background) / (maximum LDH release − medium background), where maximum LDH release was obtained from cells treated with the lysis reagent supplied with the kit.

### Immunoblotting and immunoprecipitation

For Western blotting analysis, cells were harvested and lysed in RIPA lysis buffer (Cat. P0013B, Beyotime) supplemented with complete protease inhibitor cocktail immediately before use. Cell lysates were mixed with SDS-PAGE sample loading buffer (Cat. P0015, Beyotime), boiled and separated by 12% SDS-PAGE and transferred onto polyvinylidene difluoride (PVDF) membranes (Millipore). After blocking, membranes were probed with the indicated antibodies.

For immunoprecipitation assays, cell extracts were prepared using lysis buffer containing 25 mM Tris (pH 7.4), 150 mM NaCl, 0.5% NP-40, 0.5% sodium deoxycholate and protease inhibitor cocktail. Lysates were incubated with anti-Flag affinity beads at 4℃ overnight. The beads were washed three times with the same lysis buffer, and bound proteins were eluted with SDS-PAGE sample loading buffer by boiling for 10 min.

### LPS pull-down assay

Biotinylated LPS from *E. coli* O111:B4 and biotinylated Pam3CSK4 were immobilized on Pierce™ Streptavidin beads (Cat. 20359, Thermo Fisher) by incubation for 6 h, with 1 μg of each biotinylated ligand used per reaction. The beads were washed three times with lysis buffer to remove unbound ligands. For the pull-down assays, equal amounts of lysates from transfected cells were incubated with ligand-bound beads at 4°C for 12 h. For competition assays, cell lysates were pre-incubated with 5 μg unlabeled LPS or Pam3CSK4 at 4°C for 2 h before incubation with biotinylated LPS-bound streptavidin beads. After incubation, the beads were washed with lysis buffer, and bound proteins were eluted by boiling in SDS-PAGE sample loading buffer and analyzed by immunoblotting.

### Caspase activity assays

Caspase activity was determined by measuring AMC fluorescence generated from the hydrolysis of the fluorogenic substrate Z-VAD-AMC. Briefly, 1 μg CmiCASP1 or its mutants was incubated with 1 μg of LPS or Pam3CSK4 in a 100 μL reaction buffer containing 50 mM HEPES (pH 7.2), 150 mM NaCl, 3 mM EDTA, 0.005% (v/v) Tween-20 and 10 mM DTT. After incubation for 1 h, Z-VAD-AMC was added to a final concentration of 200 μM, and the reactions were further incubated at 37°C for 30 min. Fluorescence was measured at 450 nm with excitation at 365 nm using a fluorescence multi-well plate reader (Promega).

### In vitro cleavage assays

To assess LPS-induced CmiCASP1-mediated cleavage of CmiGSDMA/B, 1 μg of CmiCASP1 or its mutants was incubated with or without 1 μg of LPS at 37°C for 1 h in a 100 μL reaction containing 50 mM HEPES (pH 7.2), 150 mM NaCl, 3 mM EDTA, 0.005% (v/v) Tween-20 and 10 mM DTT. CmiGSDMA/B (3 μg) was then added, followed by incubation at 37°C for another 1 h. The reactions were stopped by addition of SDS-PAGE sample loading buffer and boiling, and cleavage products were analyzed by immunoblotting.

### Bacterial colony-forming unit assays

Bacterial strains were streaked on appropriate agar plates and incubated overnight at 37℃. The next day, a single colony was inoculated into the corresponding fresh growth medium and cultured at 37℃ until the OD600 reached 0.5. Bacteria were then collected by centrifugation, washed with ice-cold sterile PBS, and resuspended in PBS. HEK293T cells seeded in 100-mm culture dishes were transfected with the indicated expression plasmids. At 24 h post-transfection, cells were collected and lysed in PBS by ultrasonication. Cell lysates were centrifuged at 12,000 × *g* for 10 min, and the supernatants were filtered through a 0.22 μm filter and concentrated 10-fold. Bacteria (3 μL) were mixed with 27 μL of the concentrated supernatants and incubated at 37℃ for 40 min. The reactions were serially diluted from 10^0^ to 10^5^, and 5 μL of each dilution was plated on the corresponding agar plates for CFU determination. Only dilutions yielding 30-300 colonies were used for CFU calculation.

### Bacterial live/dead assay

After incubation with concentrated supernatants at 37°C for 40 min, bacterial samples were stained with DMAO and PI for 15 min at room temperature using the LIVE/DEAD Bacterial Viability Kit (Cat. C2030S, Beyotime) according to the manufacturer’s instructions. Fluorescence images of bacteria were captured by a confocal microscope (Nikon Eclipse Ti2). Bacterial flow cytometry was performed using a FongCyte flow cytometer.

### Transcriptome analysis

Eleven RNA-seq datasets of *C. plagiosum* tissues were from the NCBI Sequence Read Archive database (SRR26468987, SRR26468991, SRR26468998, SRR26468999, SRR26469001, SRR26469005, SRR26469012, SRR46469064, SRR46469070, SRR26469092, SRR26469094). Clean reads were obtained from raw reads using fastp (v0.23.4) (Chen *et al*., 2018). The clean reads were aligned to the reference genome (GCF_004010195.1) using HISAT2 (v2.2.1) (Kim *et al*., 2019), and counts were generated with featureCounts (v2.0.6) (Liao *et al*., 2014). Counts were normalized to TPM values to compare expression across tissues. A heatmap showing the expression patterns of target genes was generated using R.

### Statistical analysis

Statistical analyses were performed with GraphPad Prism 10 software. Data are presented as mean ± SD unless otherwise indicated. Comparisons between two groups were performed using an unpaired Student’s *t* test. *p* < 0.05 was considered statistically significant.

## Data availability

All data supporting the conclusions of this study are included in the paper and its source data files. The newly generated materials in this study are available from the corresponding author upon reasonable request.

## Author contributions

Conceptualization: S.C.Y. and X.X.W.

Methodology: S.C.Y., X.X.W., and R.Z.

Investigation: X.X.W., R.Z., X.X.J., X.L.W., S.Z.L., Z.W.H., G.J.Z.

Visualization: X.X.W. and R.Z.

Writing - original draft: S.C.Y. and X.X.W.

Writing - review and editing: S.C.Y. and A.L.X.

Supervision: S.C.Y.

Funding acquisition: S.C.Y.

## Figure legends

**Figure 1—figure supplement 1. Characteristics of CmiGSDMA/B and CmiCASP1 in *Callorhinchus milii*.**

(**A**) Phylogenetic analysis of CmiGSDMA/B. GSDM members identified in *C. milii* are labeled in red. The yellow stars indicate the GSDMA/B branches of cartilaginous fish. (**B**) The domain architecture of *C. milii* caspases predicted by InterProScan, showing the presence and arrangement of N-terminal interaction modules, such as CARD and DED domains, and the conserved caspase protease core comprising the p20 and p10 domains. (**C**) A heatmap showing sequence similarity among full-length caspases. Higher sequence similarity is indicated by darker blue. CmiCASP1 is highlighted in red. (**D**) Multiple sequence alignment shows the key amino acid residues in HsCASP1 that form the substrate-binding pockets and catalytic site, as well as their corresponding positions in CmiCASP1. Sequence and structural analyses predict that the S1 pocket residues of CmiCASP1 (R173, Q277, S332) correspond to those of HsaCASP1 (R179, Q283, S339), whereas the S2, S3, and S4 pocket residues V331/F333, R334, and D335/I341/K374 correspond to V338/W340, R341, and H342/V348/R383 in HsaCASP1, respectively. The conserved catalytic motif QACRG is also indicated, with C279 in CmiCASP1 corresponding to the catalytic cysteine C285 in HsaCASP1.

**Figure 4—figure supplement 1. Homologs of inflammasome components in cartilaginous fish.**

(**A**) Schematic representation of key inflammasome components and domains architectures in *C. milii*. (**B, C**) Multiple sequence alignment of IL-1β and IL-18 homologs, respectively. The alignments were generated using the MAFFT algorithm. Conserved amino acid residues are highlighted in blue. The red box represents the predicted or previously identified caspase-1 cleavage sites. (**D**) Schematic representation of the domain architectures of inflammasome-related homologs in *C. plagiosum.* (**E**) A heatmap of inflammasome-related gene expression across eleven tissues in *C. plagiosum*. Color intensity corresponds to expression level, with darker shades indicating higher expression.

**Figure 5—figure supplement 1. Effect of N241 on the activity of *S. aureus*.**

(**A**) Representative serial dilution plating of *S. aureus* treated with the indicated GSDM lysates. *S. aureus* suspended in PBS was incubated with the indicated lysates at 37°C for 40 min, serially diluted 10-fold and plated on LB agar for CFU determination. (**B**) Representative fluorescence microscopy images of bacterial viability assessed by DMAO/PI staining. Bacteria were incubated for 40 min with the indicated concentrated lysates collected from transfected HEK293T cells. (**C**) Sequence alignment showing basic residues within the predicted basic patches (BPs) of GSDM proteins. Basic residues within the BPs are highlighted in red, and dashes indicate alignment gaps. Red labels below the alignment indicate the corresponding basic residues in CmiGSDMA/B.

## Source Data legends

**Figure 1–Source Data 1**

Source data for Annexin V^+^/PI^+^ quantification in Figure 1I.

**Figure 1–Source Data 2**

Source data for LDH release quantification in Figure 1J.

**Figure 1–Source Data 3**

Source data for cell viability quantification in Figure 1K.

**Figure 1–Source Data 4**

Original file for the western blot analysis in Figure 1L.

**Figure 2–Source Data 1**

Original file for the western blot analysis in Figure 2B.

**Figure 2–Source Data 2**

Source data for LDH release quantification in Figure 2C.

**Figure 2–Source Data 3**

Source data for Annexin V^+^/PI^+^ quantification in Figure 2D.

**Figure 2–Source Data 4**

Source data for cell viability quantification in Figure 2E.

**Figure 2–Source Data 5**

Original file for the western blot analysis in Figure 2H.

**Figure 2–Source Data 6**

Source data for LDH release quantification in Figure 2I.

**Figure 2–Source Data 7**

Source data for Annexin V^+^/PI^+^ quantification in Figure 2J.

**Figure 2–Source Data 8**

Source data for cell viability quantification in Figure 2K.

**Figure 3–Source Data 1**

Source data for Annexin V^+^/PI^+^ quantification in Figure 3C.

**Figure 3–Source Data 2**

Source data for LDH release quantification in Figure 3D.

**Figure 3–Source Data 3**

Source data for cell viability quantification in Figure 3E.

**Figure 3–Source Data 4**

Original file for the western blot analysis in Figure 3H.

**Figure 3–Source Data 5**

Source data for Annexin V^+^/PI^+^ quantification in Figure 3I.

**Figure 3–Source Data 6**

Source data for LDH release quantification in Figure 3J.

**Figure 3–Source Data 7**

Source data for cell viability quantification in Figure 3K.

**Figure 3–Source Data 8**

Source data for Annexin V^+^/PI^+^ quantification in Figure 3M.

**Figure 3–Source Data 9**

Source data for LDH release quantification in Figure 3N.

**Figure 3–Source Data 10**

Source data for cell viability quantification in Figure 3O.

**Figure 3–Source Data 11**

Original file for the western blot analysis in Figure 3P.

**Figure 4–Source Data 1**

Original file for the western blot analysis in Figure 4E.

**Figure 4–Source Data 2**

Original file for the western blot analysis in Figure 4F.

**Figure 4–Source Data 3**

Source data for fluorescence intensity quantification in the caspase activity assay shown in Figure 4G.

**Figure 4–Source Data 4**

Original file for the western blot analysis in Figure 4H.

**Figure 5–Source Data 1**

Source data for CFU quantification in bacterial killing assays shown in Figure 5A.

**Figure 5–Source Data 2**

Source data for DMAO^+^/PI^+^ quantification in *E. coli* treated with the indicated gasdermin lysates shown in Figure 5D.

**Figure 5–Source Data 3**

Source data for DMAO^+^/PI^+^ quantification in *S. aureus* treated with the indicated gasdermin lysates shown in Figure 5E.

**Figure 5–Source Data 4**

Source data for CFU quantification in bacterial killing assays shown in Figure 5I.

**Figure 5–Source Data 5**

Source data for LDH release quantification in Figure 5K.

## Supplementary file legends

**Supplementary file 1**

Species, gene names and accession numbers of caspase sequences used for phylogenetic analysis in Figure 1C.

